# Injectable Nanoelectrodes Enable Wireless Deep Brain Stimulation of Native Tissue in Freely Moving Mice

**DOI:** 10.1101/2020.03.14.978676

**Authors:** Kristen L. Kozielski, Ali Jahanshahi, Hunter B. Gilbert, Yan Yu, Önder Erin, David Francisco, Faisal Alosaimi, Yasin Temel, Metin Sitti

## Abstract

Devices that electrically modulate the central nervous system have enabled important breakthroughs in the management of neurological and psychiatric disorders. Such devices typically have centimeter-scale dimensions, requiring surgical implantation and wired-in powering. Using smaller, remotely powered materials could lead to less invasive neuromodulation. Herein, we present injectable magnetoelectric nanoelectrodes that wirelessly transmit electrical signals to the brain in response to an external magnetic field. Importantly, this mechanism of modulation requires no genetic modification of the brain, and allows animals to freely move during stimulation. Using these nanoelectrodes, we demonstrate neuronal modulation *in vitro* and in deep brain targets *in vivo*. We also show that local thalamic modulation promotes modulation in other regions connected via basal ganglia circuitry, leading to behavioral changes in mice. Magnetoelectric materials present a versatile platform technology for less invasive, deep brain neuromodulation.

## Introduction

Electrical communication with and modulation of the central nervous system (CNS) are essential to our current understanding of neurobiology, and in the diagnosis and treatment of neurological disorders. Using sensing and/or modulation of neural electrical activity, key therapeutic CNS interventions have allowed remarkable medical breakthroughs. For more than 30 years, deep brain stimulation (DBS) has provided patients with symptom relief from Parkinson’s Disease, as well as other disorders, using electrodes wired into deep targets within the brain (1). More recently, closed-loop control of epidural electrical stimulation enabled walking in patients with spinal cord injury (2). Importantly, such devices function in freely moving patients, enabling daily activity and chronic patient use.

In recent years, efforts to make neural intervention less invasive, longer-lasting, and safer have progressed the capabilities of neural devices (for review, see (3)). A key challenge of such devices is powering, and wired-in powering can require that patients undergo surgical battery changes, every 3-5 years in the case of DBS devices (4). Instead, neural devices that are remotely powered have emerged using magnetic induction (5), opto-electric signaling (6-8), acoustic powering of piezoelectric materials (9-14), magnetic heating (15), piezoelectric powering of LEDs (16, 17), or magnetoelectric materials (18), instead of a wired-in battery.

Like conventional DBS electrodes, centimeter-scale devices require surgery and implantation of hardware external to the CNS, which risks brain hemorrhage, infection, and damage during daily activity (4). Thus, several neural device technologies have instead turned to smaller (nano-to millimeter-scale) devices, which can be completely implanted within the CNS, potentially via injection.

However, smaller size can make powering of neural devices more difficult. Remotely powered devices using magnetic induction (5), or opto-electric signaling (6, 7) thus far are limited in their tissue penetration depth, maximally reaching 1 cm and 6 mm, respectively (19). Ultrasound-powered piezoelectric devices are perhaps the most promising of these technologies, recently showing recording at multiple sites through 5 cm of tissue phantom material with a sub-mm^3^ device (10). Modulation with piezoelectric devices, however, has currently only been demonstrated in the peripheral nervous system using millimeter-scale devices, or *in vitro* (12-14). As power transmission is typically done at the mechanical resonance frequency of such devices, this creates a fundamental tradeoff where an increasingly smaller device with a higher resonance frequency can be powered at increasingly shallower tissue depths (20, 21). Thus, resonant frequency signaling creates an obstacle to modulating deep brain targets with an injectable-sized device.

To circumvent signal transmission challenges, other strategies have used genetic neuronal modification and magnetic nanoparticles (15), or piezoelectrically-powered LEDs (16, 17) to trigger ion channel opening. However, the dependence of such technologies on genetic tissue modification creates regulatory barriers to their translation into patients. Wireless modulation of neural activity is clinically available using transcranial magnetic stimulation (TMS), which requires no implanted device (22). However, TMS only modulates cortical tissue (23), and has a depth-focal area tradeoff (24, 25), making DBS via TMS currently impossible.

To achieve wireless signal transmission to injectable devices, we have used magnetoelectric nanoelectrodes, which couple magnetic and electric signals (**Fig. 1A**). Technologies using magnetoelectric materials for neuromodulation have previously been explored. A centimeter-scale device has been used for DBS (18), and in other work, magnetoelectric nanoparticles were used but not reported to modulate activity in the deep brain (26). However, both demonstrate the promise of magnetoelectric materials for neural devices.

**Figure 1.**
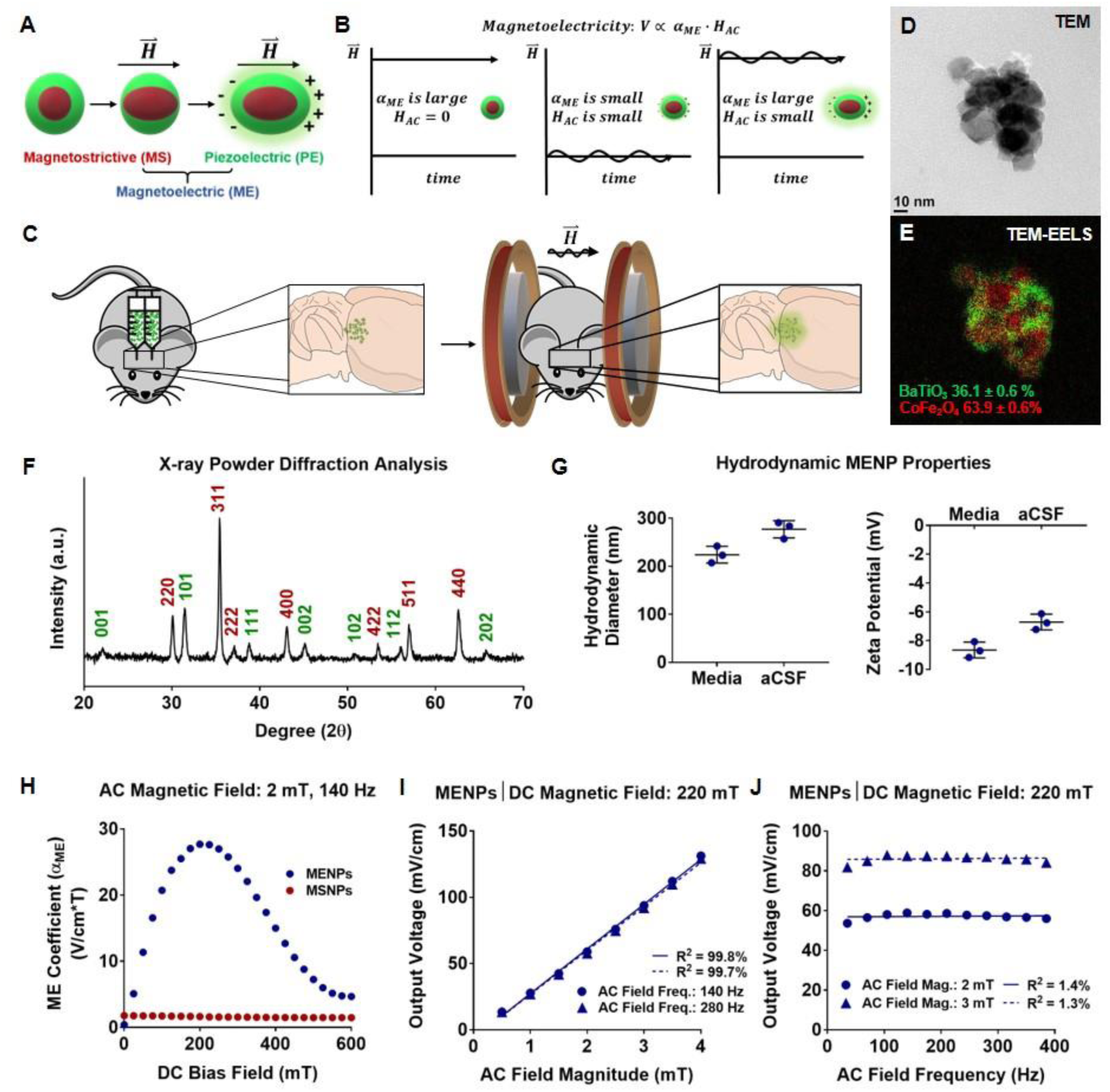
Material and magnetoelectric characterization of MENPs made from magnetostrictive and piezoelectric phases demonstrates wireless electric field generation. Schematic demonstrating two-phase magnetoelectricity in materials made from magnetostrictive and piezoelectric materials that are strain-coupled (A). Schematic demonstrating the rationale for using a large DC magnetic field overlaid with an AC field to generate optimal magnetoelectric output (B). Diagram of method of *in vivo* MENP administration. MENPs are injected bilaterally into the thalamic region of mice, and MENPs are wirelessly stimulated using an AC and DC magnetic field (C). Transmission electron microscope (TEM) (D) and TEM-electron energy loss spectroscopy (TEM-EELS) images (E) show MENP morphology and BaTiO_3_/CoFe_2_O_4_ phases (green and red, respectively), with quantitative elemental analysis measurement of the molar percentage of each material (E). MENPs were analyzed via X-ray powder diffraction (XRD) to confirm the perovskite crystal structure of BaTiO_3_ (green) and the spinel crystal structure of CoFe_2_O_4_ (red) (F). Dynamic light scattering (DLS) was used to characterize MENP hydrodynamic properties in cell culture media and aCSF (G). The input-output magnetoelectric coefficient was measured as a function of DC bias field in MENPs and MSNPs (H). Output voltage of MENPs was measured using a 220 mT DC field, while varying AC field magnitude (I) or AC field frequency (J). Plots show individual points with mean ± SD (n = 3) (G), and individual points fitted to a linear correlation (I,J).

Herein, we report wireless DBS *in vivo* using injectable, magnetoelectric nanoelectrodes. They are implanted into the subthalamic area via stereotactic infusion, and stimulated using an external magnetic field at non-resonant frequencies, and in freely moving mice (**Fig. 1C**).

In particular, we made two-phase magnetoelectric nanoparticles (MENPs) using magnetostrictive CoFe_2_O_4_ nanoparticles (MSNPs) coated with piezoelectric BaTiO_3_. The two materials are strain-coupled via sol-gel growth of BaTiO_3_ on CoFe_2_O_4_ nanoparticles. Wireless particle stimulation is achieved by application of a magnetic field, which creates strain in CoFe_2_O_4_, resulting in applied strain to BaTiO_3_, thereby creating a charge separation (**Fig. 1A**). Below, we demonstrate wireless generation of an electric field across MENPs using an applied magnetic field. We then show that magnetic stimulation of MENPs enables wireless modulation of neuronal activity *in vitro* and *in vivo*. Finally, we demonstrate the therapeutic potential of this technology through its ability to modulate activity in the motor cortex and nonmotor thalamus, and to alter animal behavior.

## Results and Discussion

Two-phase MENPs were synthesized using a protocol similar to Corral-Flores *et al.* (27). The nanoparticles were characterized for morphology (**Fig. 1D,E**), magnetostrictive to piezoelectric material ratio (**Fig. 1E**), and crystal structure (**Fig. 1F**). We observed two-phase MENPs containing 36.1 ± 0.6 % BaTiO_3_ and 63.9 ± 0.6% CoFe_2_O_4_, in their perovskite and spinel crystal structures, respectively. MENP hydrodynamic properties were also characterized via dynamic light scattering (DLS) in cell culture medium and an artificial cerebrospinal fluid (aCSF) solution. Average particle diameter was measured as 224 ± 17 nm and 277 ± 18 nm, and zeta potential was measured to be −8.6 ± 0.5 mV and −6.7 ± 0.5 mV, in medium and aCSF, respectively (**Fig. 1G**).

We next measured the electrical output of MENPs under an applied magnetic field to characterize their ability to wirelessly generate an electric field. MENPs were measured as a sintered, poled pellet by attaching electrodes and measuring the output voltage via a lock-in amplifier (**Fig. S1**). A pellet containing only MSNPs was used as a negative control. To optimize our ME output, we applied a small AC magnetic field with a larger DC bias field along the same axis (**Fig. 1B**). The magnetoelectric coefficient (*α*_*ME*_), which quantifies the relationship between the input AC magnetic field and output voltage, varied nonlinearly with the DC field, as has previously been reported (28). The *α*_*ME*_ reached a maximum of 27.6 mV cm^-1^ T^-1^ at 200 and 225 mT in the MENP pellet, while the MSNP *α*_*ME*_ showed no dependence on the DC field (**Fig. 1H**). Using a DC field within the maximum *α*_*ME*_ range (220 mT), we measured a linear relationship between the AC field magnitude and the output voltage (R^2^ = 99.8% and 99.7% at AC frequencies of 140 Hz and 280 Hz, respectively) (**Fig. 1I**), which is also typical of magnetoelectric materials (28).

Importantly, we found a low correlation (R^2^ = 1.4% and 1.3% for AC magnitude 2 mT and 3 mT, respectively) between the output voltage relative to AC field frequency across the range tested (35 – 385 Hz), which covers the range of DBS frequencies found to have clinical effect (reviewed in (29)) (**Fig. 1J**). This frequency range also has little attenuation in tissue, thus improving potential signal penetration depth (20, 21).

The effect of wireless MENP signaling on neuronal cell activity was examined *in vitro* in real-time using intracellular Ca^2+^ signaling in differentiated human SH-SY5Y cells. MENPs were administered at 100 µg/mL as a suspension in the imaging medium 20 min prior to testing, using no NPs, MSNPs, and piezoelectric nanoparticles (PENPs) as controls. Prior to choosing a concentration, the toxicity of MENPs was assessed with a lactate dehydrogenase (LDH) assay and a metabolic activity assay (MTS) (**Fig. S2**). Magnetic stimulation parameters were either no field, a 225 mT (within the maximum *α*_*ME*_ range) DC field, a 6 mT, 140 Hz AC field, or both DC and AC fields together using a custom coil system (**Fig. S3**). The DC or AC magnetic fields alone were not expected to output a magnetoelectric effect sufficient to modulate neuronal activity, and were therefore used as controls. We found a significant increase in the percentage of cells exhibiting Ca^2+^ transients when MENPs were stimulated with a simultaneous AC and DC magnetic field (20.1 ± 2.3%) versus basal activity (2.8 ± 2.6%) (**Fig. 2A-C, Movie S1**). This increase was not observed when cells were exposed to the AC and DC magnetic stimulation either alone (1.0 ± 1.7%), with MSNPs (1.4 ± 1.3%), or PENPs (1.4 ± 1.2%), which supports our hypothesis that the measured increase in activity was due to magnetoelectric voltage generation. While the MENPs seem to have some effect on neuronal activity (7.2 ± 5.0%, 5.2 ± 6.0%, or 3.8 ± 5.0%, with no field, AC field only, or DC field only, respectively), this effect was not significantly different than any other negative control group (**Fig. B**,**C; Table S1, Movie S1**).

**Figure 2.**
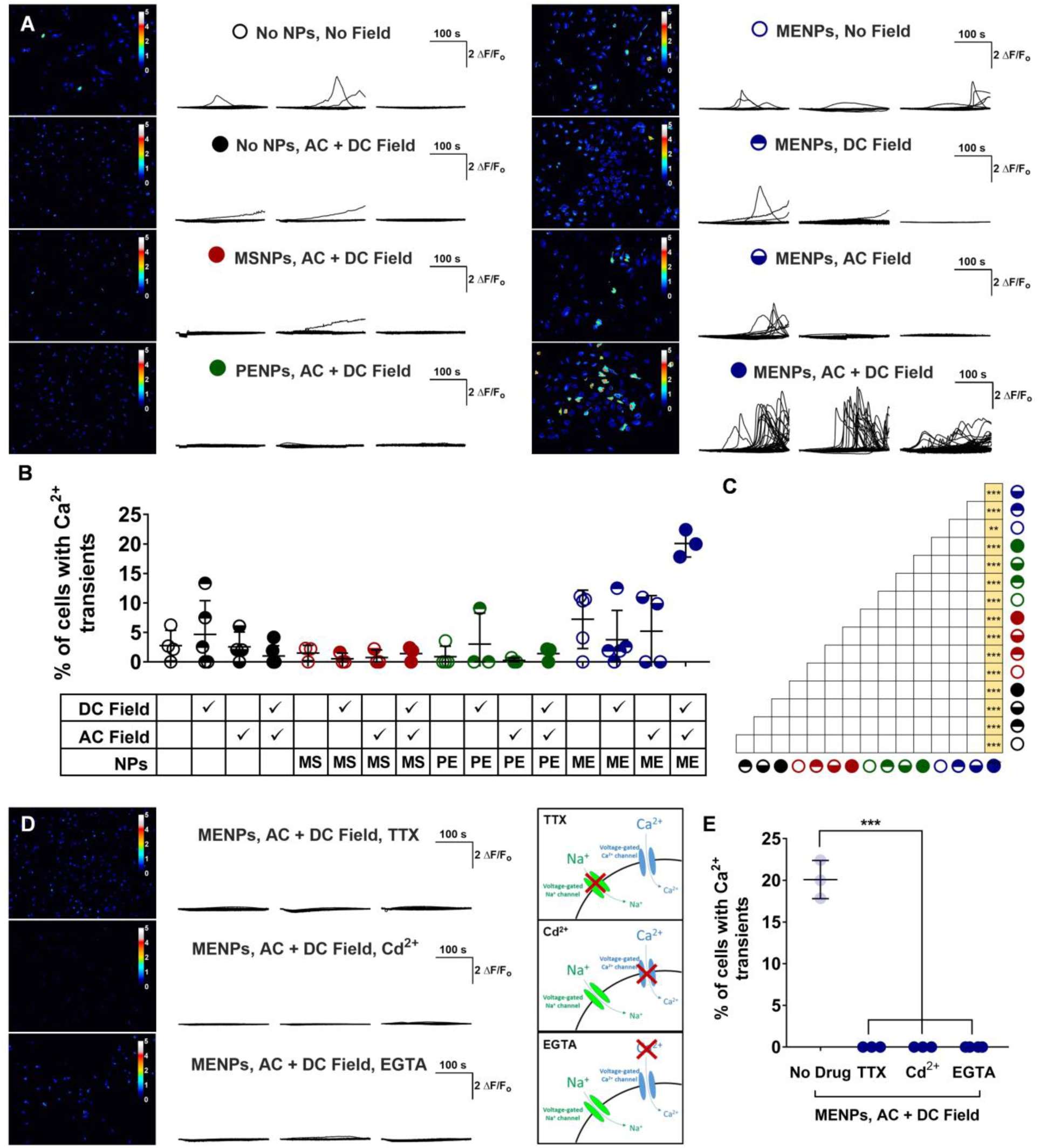
Magnetic stimulation of MENPs modulates neuronal cell activity *in vitro*. Cells were treated with MENPs, using no NPs, MSNPs, or PENPs as controls, prior to magnetic stimulation. Magnetic stimulation was either 220 mT DC (DC), 6 mT and 140 Hz AC (AC), or both DC and AC fields along the same axis (AC + DC). Neuronal activity was measured in real time via intracellular Ca^2+^ imaging using Fluo4 dye, and cell fluorescence was traced over time per cell. Images of total Ca^2+^ activity over time is shown for selected experimental groups. Calibration bars represent ΔF/F_o_ (A) The percent of cells demonstrating intracellular Ca^2+^ transients in (A) is summarized in (B) and significantly different group comparisons are marked in yellow within (C). Cells were treated with TTX, Cd^2+^, or EGTA prior to treatment with MENPs and AC and DC magnetic stimulation, and total Ca^2+^ activity over time is shown next to a diagram depicting the inhibiting activity of each drug (D). Measured Ca^2+^ transients of drug-treated cells are summarized with no drug, MENP, AC + DC Field treated cells shown as faded plot points for comparison (E). Plots show traces of Ca^2+^ activity over time in individual cells (A,D) and individual points with bars showing mean ± SD (B,E), (n = 3 – 6); ANOVA with Tukey’s post-test (C), or Dunnett’s post-test with no drug as the control (E); **p < 0.01, ***p < 0.001, unlabeled group comparisons are not significantly different.

In order to support our hypothesis that the Ca^2+^ activity we measured was related to electrophysiological cell activity, we stimulated the MENPs with AC and DC magnetic fields, but first treated the cells with either a voltage-gated Na^+^ channel blocker (tetradotoxin, TTX), a voltage-gated Ca^2+^ channel blocker (Cd^2+^), or an extracellular Ca^2+^ chelator (ethylene glycol-bis(β-aminoethyl ether)-N,N,N′,N′-tetraacetic acid, EGTA) (schematic in **Fig. 2D** showing drug activity). In the presence of each drug, the cells failed to produce any Ca^2+^ transients (**Fig. 2D**,**E**). This substantiates the dependence of our measured Ca^2+^ transients on voltage-gated ion channels and extracellular Ca^2+^ sources, supporting the relationship between our measured Ca^2+^ activity and cell electrophysiological activity.

We then sought to assess the feasibility of MENP-based neuromodulation *in vivo*. MENPs were bilaterally injected into the ventral thalamic region of naïve mice (C57Bl/6J) at a dose of 100 µg/animal, which was found to be tolerable in a dose-toxicity assessment (**Fig. S4**). This region of the brain was selected as the basal ganglia and thalamus are the most common target areas for DBS (30). Moreover, these areas involve well understood brain circuits in the fields of DBS and neuromodulation for neurological disorders. In addition to the classical basal ganglia model, new models show that parallel circuits also engage associative and limbic regions (31, 32). Therefore, the cortico-basal ganglia-thalamo-cortical circuit provides a tool to reliably investigate the effects of neuromodulation on wide range of behavioral functions.

During magnetic stimulation, mice were awake and unrestrained within our *in vivo* magnetic coil device (**Fig. S5**). As a control group, mice were treated with MENPs and a DC magnetic field only, meaning they were placed into the magnetic device, but with the AC coil remaining off. We first assessed changes in local neural activity by immunohistochemically measuring the expression of *c-fos* protein, a widely used cell activity marker (33). We found significantly more *c-fos* positive cells in the region of nanoparticle injection when animals were treated with MENPs and an AC and DC field (38.5 ± 8.0 cells), versus only a DC field (4.25 ± 3.0 cells) (**Fig. 3A-C**). This data supports our hypothesis that we could wirelessly modulate local brain activity using the magnetoelectric response of MENPs to magnetic stimulation.

**Figure 3.**
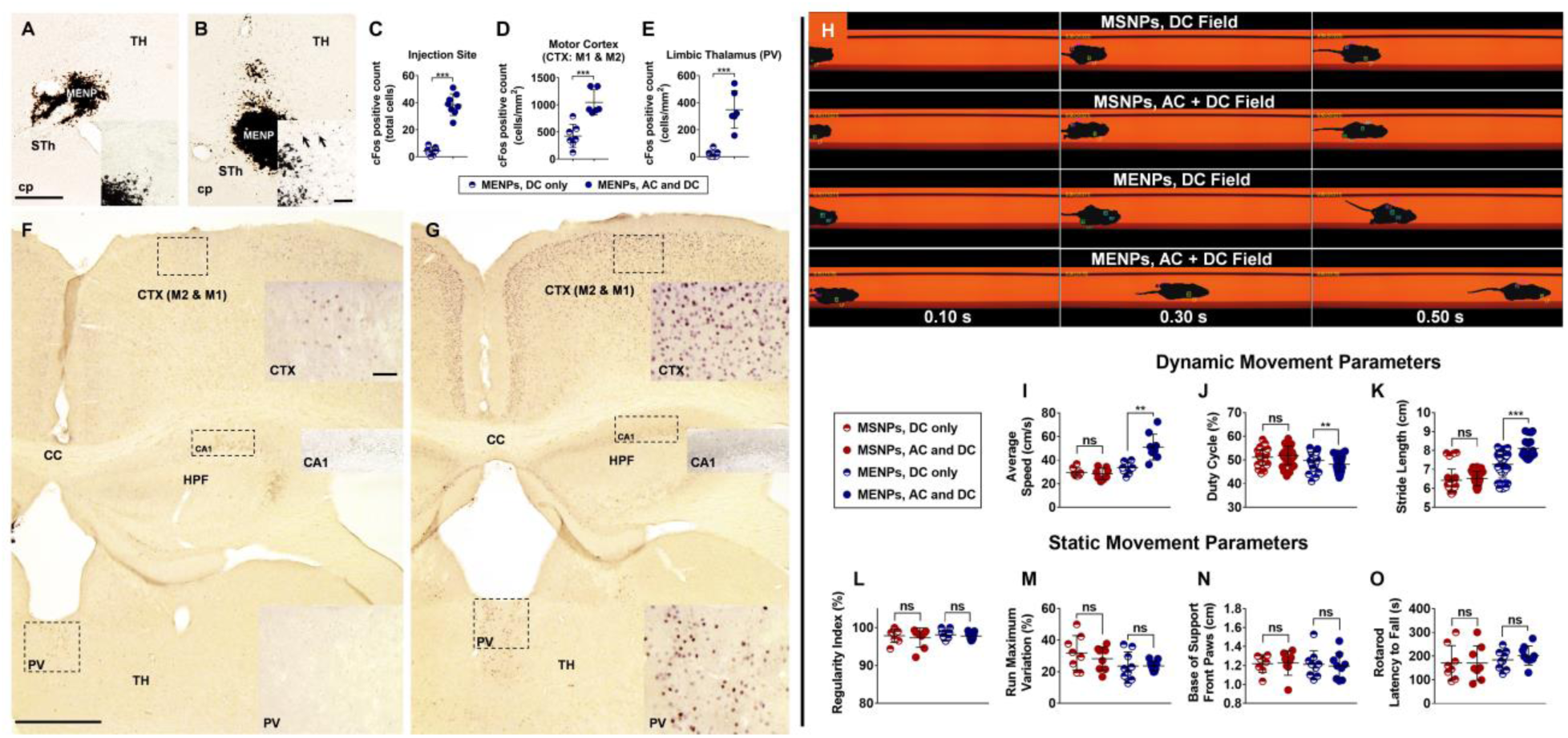
Magnetic stimulation of MENPs locally modulates neural activity in mice, yielding modulation of basal ganglia circuitry and behavioral change. Staining for *c-fos* protein locally to the MENP injection site following DC magnetic stimulation (A) or AC and DC magnetic stimulation (B) shows increased *c-fos* expression in the latter (C). Quantification of *c-fos* expression in the motor cortex (D) and limbic thalamus (E) shows increased expression when MENPs were stimulated with an AC and DC magnetic field (G) versus only a DC magnetic field (F). Time-lapse images showing mouse movement in a catwalk video recording system (H). Dynamic movement parameters as measured by the catwalk recording showed significant changes in mouse speed (I), limb duty cycle (J), and limb stride length (K) in MENP-treated mice following AC and DC stimulation versus DC stimulation, while MSNP-treated mice showed no change. Static movement parameters of mouse movement such as regularity index (L), run maximum variation (M), and front-paw base of support (N) as measured by catwalk recording did not significantly change with AC and DC versus DC only magnetic stimulation in either nanoparticle group. Rotarod latency to fall also did not significantly change with AC and DC versus DC only magnetic stimulation in either nanoparticle group (O). Scale bar, 250 µm (overview) and 50 µm (inset) (A, B, F, G). Plots show individual points with bars showing mean ± SD (C-E, I-O), (n = 6-9 mice, individual limb values for J,K); Unpaired t-test (C-E), or paired t-test (I-O), ns = not significant, **p < 0.01, ***p < 0.001.

We next wanted to determine if local thalamic neuromodulation induced by MENPs was sufficient to cause modulation in other regions of the cortico-basal ganglia-thalamo-cortical circuit. We found *c-fos* protein expression significantly higher in the motor cortex and nonmotor thalamus following stimulation with MENPs and an AC/DC magnetic field (1046.4 ± 232.4, 348.4 ± 137.7 cells/mm^2^, respectively) versus only a DC magnetic field (424.8 ± 214.9, 19.9 ± 27.6 cells/mm^2^, respectively) (**Fig. 3D-G**). Importantly, we did not observe a global change in *c-fos* protein expression, such as in the CA1 region of the hippocampus (**Fig. 3F,G**). Together, these data support our hypothesis that the measured increases in *c-fos* protein expression were due to local thalamic stimulation of the cortico-basal ganglia-thalamo-cortical circuit, and not a nonspecific, global modulation of neural activity via the magnetic field.

To determine if the induced neuromodulation would affect animal behavior, we tested mice via a Rotarod test and an automated CatWalk XT gait analysis system. Mice were injected with MENPs or MSNPs as a control. Behavior with AC and DC magnetic stimulation versus behavior with only DC magnetic stimulation was compared for each mouse. Gait and balance-related static parameters during the CatWalk test, such as regularity index, run maximum variation, and base of support, showed no significant difference following AC and DC stimulation in either nanoparticle group (MSNPs, 97.3 ± 2.5 vs. 97.9 ± 1.8%, 28.3 ± 7.6 vs. 31.9 ± 11.0%, 1.2 ± 0.1 vs. 1.2 ± 0.1 cm, DC vs. AC and DC stimulation, respectively) (MENPs, 97.8 ± 1.0 vs. 98.1 ± 1.3%, 23.7 ± 3.3 vs. 23.6 ± 8.9%, 1.2 ± 0.1 vs. 1.2 ± 0.1 cm, DC vs. AC and DC stimulation, respectively) (**Fig. 3L-N, Movie S2**). Rotarod testing also showed no significant difference in latency to fall with either nanoparticle group (MSNPs, 170.4 ± 72.9 vs. 170.9 ± 73.2 s; MENPs, 202.6 ± 40.4 vs. 184.0 ± 40.5 s, DC vs. AC and DC stimulation,) (**Fig. 3O**). While we anticipated no improvement in motor function, as we tested only naïve mice, these results are important to demonstrate that we saw no detrimental effect to the gait and balance of the animals due to neuromodulation via MENPs. This finding of no generalized behavioral change also corresponds to our *c-fos* expression findings, in which we found only selective expression changes.

Conversely, in analyzing the dynamic parameters of the catwalk test, which are indicative of animal speed, we found a significant difference in the behavior of MENP-treated animals that was not observed in MSNP-treated animals (**Fig. 3I-K, Movie S2**). The average speed, duty cycle of each limb, and stride length of each limb all changed significantly in MENP-treated mice following AC and DC stimulation (51.1 ± 10.9 vs. 33.6 ± 4.8 cm/s, 48.1 ± 3.0 vs. 49.9 ± 3.5%, and 8.1 ± 0.5 vs. 7.3 ± 0.7 cm, DC vs. AC and DC stimulation, respectively), but not in MSNP-treated mice (28.3 ± 5.0 vs. 29.4 ± 3.8 cm/s, 51.9 ± 3.9 vs. 51.1 ± 3.2%, 6.5 ± 0.4 vs. 6.4 ± 0.6 cm, DC vs. AC and DC stimulation, respectively). As anxiety can be induced via stimulation of the thalamus, we believe that the measured changes in animal speed are due to induced anxiety via selective modulation of the thalamo-cortical circuit. The combined results of *c-fos* protein immunohistochemistry and animal behavioral tests support the conclusion that magnetically-stimulated MENPs wirelessly modulated basal ganglia circuitry to affect brain activity and animal behavior.

## Conclusion

This work demonstrates the potential of magnetoelectric materials as nanoelectrodes for wireless electrical modulation of deep brain targets. Herein, we have shown that we can stimulate MENPs with a magnetic field to remotely generate electric polarization of the MENPs. We have shown evidence that non-resonant frequency magnetic stimulation of MENPs locally modulates neuronal activity *in vitro* and *in vivo*. We have also demonstrated that this modulation is sufficient to change animal behavior, and to modulate other regions of the cortico-basal ganglia-thalamo-cortical circuit. Future work will be key to optimizing magnetoelectricity based neural devices and understanding the abilities and limitations of this technology. Magnetoelectric nanoelectrodes show promise for new technologies in wireless neural devices.

## Materials and Methods

### Magnetoelectric nanoparticle (MENP) synthesis

MENPs were synthesized in a manner similar to Corral-Flores *et al. (29)*. CoFe_2_O_4_ nanoparticles (30 nm, Sigma-Aldrich) were suspended in dH_2_O at a concentration of 10 mg/mL to 80°C while stirring. Oleic acid was added to the suspension at 30 wt.% with respect to CoFe_2_O_4_, the temperature was raised to 90°C for 30 min, then lowered to 60°C. Octane was added to the suspension at a 1:1 ratio to the dH_2_O volume, which separated oleic acid coated CoFe_2_O_4_ particles into the organic layer. The organic layer was then washed with dH_2_O three times. Barium acetate (BaAc) and titanium butoxide (TiButO) were dissolved in glacial acetic acid with stearic acid (final concentration 0.01%) such that the final molar ratio of BaTiO_3_ to CoFe_2_O_4_ was 1:3. This solution was stirred and heated to 90°C, the CoFe_2_O_4_ solution was added, as well as 2-methoxyethanol at a final volume concentration of 30%. The solution was dried, calcined at 700°C for 2 h, and then ground with a mortar and pestle. To select for particles with better colloidal stability, MENPs were suspended in dH_2_O, centrifuged for 1 min at 10 G, and particles within the supernatant were kept for further experiments.

### X-ray diffraction (XRD) analysis of MENP crystal structure

XRD analysis of MENPs was carried out on a Bruker D8 Advance Powder Diffractometer using Cu radiation generated at 40 kV/40 mA with a Bragg-Brentano beam path. A divergence slit at 0.5°, anti-scatter slits at 2° and 4°, and Soller slits were used. The output beams were received using a VÅNTEC-1 1-dimensional detector. Peaks were identified using the International Centre for Diffraction Data (ICDD) database. MathWorks® MATLAB software was used to baseline-correct the spectrum, using the *msbackadj* function.

### Elemental Analysis of MENPs to determine chemical composition

MENP elemental analysis was carried out via Inductively Coupled Plasma – Optical Emission Spectrometry (ICP-OES) using a Spectro Ciros spectrometer (Kleve, Germany). MENPs were first dissolved in an aqueous solution of 3% HNO_3_ and 1% HF prior to sample loading in the spectrometer. Data were analyzed using Spectro ICP Analyzer software to detect Ba, Ti, Co and Fe spectra. Data are presented as the mean ± SD of each element measured within BaTiO_3_ and CoFe_2_O_4_.

### Transmission electron microscopy (TEM) and electron energy loss spectroscopy (EELS) analysis of MENP morphology

MENPs were prepared for TEM analysis by drop-casting an aqueous suspension onto C-coated-Cu TEM grids and air-drying. TEM and TEM-EELS images were acquired using a ZEISS Sub-Electron-volt Sub-Angstrom Microscope (SESAM). Data was acquired in TEM mode at 200 kV. For EELS, we acquired energy-filtered TEM (EFTEM) spectrum images from 30 to 120 eV, with 3 eV steps and 4X binning. After data acquisition, the EELS signal from Ba (N_4,5_ edge, 90 eV) and Fe (M_2,3_ edge, 54 eV) was extracted and used for the elemental map.

### Analysis of MENP hydrodynamic properties

Hydrodynamic diameter and zeta-potential of MENPs were measured via dynamic light scattering (DLS) using a Wyatt Mobius™ DLS Instrument and analyzed via Wyatt DYNAMICS software. MENPs were diluted to a concentration of 100 µg/mL in either our cell culture differentiation media (see below) or an artificial cerebrospinal fluid (aCSF) solution (34) during the measurements. Data was analyzed from three independent experiments.

### Formation of sintered pellets of nanoparticles and pellet wiring

For ME measurement of pellets, 0.65 g of MENPs were mechanically pressed into a pellet of diameter 8 mm using 6 tonnes/cm^2^ of pressure, then sintered at 1150°C for 12 h. MSNP pellets were prepared in the same way but using only CoFe_2_O_4_ nanoparticles. The circular surfaces of the pellets were painted with conductive silver glue to attach copper plates (**Fig. S1A**). Pellets were heated to 140°C, electrically poled at 1 kV/mm thickness for 5 min, then allowed to cool to room temperature while maintaining the applied voltage. The pellets were then wired to a charge amplifier. The pellet and charge amplifier were enclosed in a Faraday shield, and connected externally to a lock-in amplifier for voltage measurement (**Fig. S1B-H**).

### Design of charge amplifier

For electrical measurement of the magnetoelectric response of pellets, a charge amplifier is used to eliminate the effects of stray capacitance on the measurement of the piezoelectric charge. The battery-powered amplifier was constructed on a standard FR4 printed circuit board, which was placed within the Faraday. The charge amplifier uses an operational amplifier circuit (**Fig. S1F-H**) based on the Texas Instruments OPA340. The amplifier has a high-pass characteristic with a –3 dB frequency of 3 Hz, and the calculated gain of the circuit in the passband is 200 mV/pC.

### Magnetoelectricity measurements

A Microsense EZ vibrating sample magnetometer (VSM) was used as a DC magnetic field source, and was modified to hold an additional, smaller Helmholtz coil. This was powered with a signal generator (35 – 385 Hz sine wave) connected to a linear voltage amplifier (Hewlett Packard) to provide current to the smaller coils, generating an AC magnetic field in the plane of the sample. The pellet was oriented such that the AC and DC magnetic fields were parallel to the pellet’s central axis (**Fig. S1E**). The AC magnetic field magnitude was measured using a gaussmeter prior to experimentation. Pellets were demagnetized prior to all measurements.

### Culture and differentiation of SH-SY5Y neuronal cells

SH-SY5Y cells were purchased from DSMZ (ATCC® CRL-2266™). Maintenance cultures were grown in DMEM/F12 (Gibco) with 10% fetal bovine serum (FBS) and 1% penicillin/streptomycin, at 37°C with 5.0% CO_2_. Media was changed every 3 to 4 days. Prior to plating for experiments, wells were coated with 5 µg/mL laminin in phosphate-buffered saline (PBS) with Ca^2+^/Mg^2+^ for 1 h at 37°C. For Ca^2+^ signaling experiments, cells were plated at a concentration of 20,000/cm^2^ onto cell culture treated, 4-well IBIDI® µ-slides. For toxicity analysis, cells were plated at a concentration of 20,000/cm^2^ onto cell culture treated 96-well plates. Experimental cultures were differentiated in DMEM/F12 medium containing 1% FBS, 1% penicillin/streptomycin, and 10 µM retinoic acid (Sigma-Aldrich) for 4 days prior to all experiments.

### Analysis of cell toxicity

MENPs were suspended in experimental cell culture medium at a concentration of 0, 50, 100, 200, or 300 µg/mL and added to cells. Toxicity was assessed at 24 h following MENP administration via a CyQUANT™ lactate dehydrogenase (LDH) assay kit (ThermoFisher Scientific), as well as a CellTiter 96® AQueous One MTS assay. Assay results were read using a BioTek® Synergy™ 2 Microplate Reader (**Fig. S2**). Each experiment was tested within 4 wells, and the average of these values was recorded to provide a single data point. The data was analyzed from three independent experiments.

### *In vitro* magnetic stimulation

A magnetic stimulation setup was designed to fit into a Zeiss Axio Observer A1 microscope, and to hold a 4-well IBIDI® µ-slide (**Fig. S3**). A DC magnetic field was provided by three permanent NdFeB magnets (N42, 6 cm diameter, 5 mm height; Supermagnete) on either side of the cells to generate a 225 mT field at the center of the cell culture well. A magnetic coil was used to provide an AC magnetic field along the same axis. AC signals were generated by a National Instruments™ DAQ USB X-Series device, controlled via LabVIEW software, and amplified by a class D audio amplifier. For all experiments with AC magnetic stimulation, the AC field component was a 6 mT sine wave at 140 Hz applied during the time window of 10 – 30 s during the time lapse recording. AC and DC magnetic field magnitudes were verified with a magnetometer.

### Ca^2+^ signaling experiments

Cells were loaded with 1 µM Fluo4-AM dye (ThermoFisher Scientific) in Live Cell Imaging Solution (LCIS, Invitrogen) for 30 min at 37°C. Experimental suspensions of no NPs, MENPs, PENPs, or MSNPs were prepared at 100 µg/mL in LCIS. After Fluo4 loading, cells were washed 3X with LCIS, and particle suspension solutions were added. Cells with particles were incubated for 20 min at 37°C to allow Fluo4 to de-esterify, then moved onto a Zeiss Axio Observer A1 microscope mounted with the *in vitro* coil system. For experiments using inhibitory drugs, Fluo4 loading was carried out as described above, and drugs were added in the LCIS solution with MENPs after washing. For EGTA, PBS was used instead of LCIS, and was added during the Fluo4 loading step. TTX was added at a concentration of 100 nM, CdCl_2_ (Cd^2+^) was added at 100 µM, and EGTA at 5 mM, which have previously been determined to be inhibitory but nontoxic concentrations (14).

Fluo4 was excited using a 470 nm LED with a 484/25 nm excitation filter, and observed through a 519/30 nm emission filter. Time lapse images were taken at 10X magnification, every 1 s for 240 s using 50 ms illumination, and recorded using a Zeiss Axiocam 503 mono camera (2.8 megapixels). Data was collected from 3 – 6 independent experiments per group.

Time lapse recordings were analyzed using ImageJ software. Briefly, the first 10 images of each time lapse were stacked into a single image to enable region of interest (ROI) selection. Following brightness normalization, blurring, background subtraction, and thresholding, ROIs were selected from this image using the Analyze Particles function (with all settings remaining consistent for all time lapses). These ROIs were then overlaid onto the completely unmodified time lapse series, and the mean gray value within each ROI was recorded for each frame. These values were then used to calculate Ca^2+^ transient amplitudes as ΔF/F_o_. Cells positively showing Ca^2+^ transients were calculated using MathWorks® MATLAB software, using a linear baseline correction and the *peakfinder* function. Images in **Fig. 2A,D** were generated by creating a maximum value Z-stack of the entire video.

### Animals

Experiments were performed on 68 male naïve mice (C57Bl/6J; Jackson Laboratory). Mice were socially housed under controlled conditions (21±2°C and 40-60% humidity) in a reversed 12h day/night cycle (lights on, 7 p.m.) until they had received surgery. Mice were given ad libitum access to food and water. At the time of surgery, mice were 3 months of age. Experiments were conducted according to the directive 2010/63/EU for animal experiments and in agreement with the Animal Experiments and Ethics Committee of the Maastricht University, Maastricht, The Netherlands.

### Stereotactic nanoparticle administration

Buprenorphine 0.1 mg/kg was subcutaneously injected half an hour prior to surgery as an analgesic. Inhalational anesthesia was induced and maintained with isoflurane (Abbot Laboratories, Maidenhead, UK) at 4% and 1.5-3%, respectively. After adequate induction of the anesthesia, the mouse was placed in a small animal stereotaxic frame (Stoelting, Dublin, Ireland) and fixed by ear bars with zygoma ear cups (Kopf, Los Angeles, United States of America) and a mouse gas anesthesia head holder (Stoelting, Dublin, Ireland). To maintain body temperature at 37°C throughout the whole procedure, the mouse was placed on a thermo-regulator pad. An ocular lubricant was applied to prevent drying of the eyes. A subcutaneous injection of Lidocaine 1% (Streuli Pharma, Uznach, Switzerland) at the incision side was given for local anesthesia.

Consecutively, burr holes above the subthalamic area (AP: −2.06 mm, ML: −1.50 mm, DV: −4.50) was made and a total of 2 µl of MENPs or MSNPs were injected with a microinjection apparatus Nanoject II (Drummond Scientific). In phase-I in vivo experiment MENPs injection was conducted only in right hemisphere to compare microglia and astrocytes population between injected and intact hemispheres. The infusion rate was 100 nl/min. After the injections, the syringe needle remained inside the brain for another 10 min prior to a slow withdrawal.

### *In vivo* magnetic stimulation

All *in vivo* magnetic stimulation was carried out using a custom coil system that would allow mice to move freely during the experiments. The animal experiment setup was designed to provide a 220 mT DC magnetic field with a 6 mT, 140 Hz AC magnetic field along the same axis at the center of the animal chamber. Images and the design of the *in vivo* coil system are shown in **Figure S5**. The structure was 3D-printed with Acrylonitrile Butadiene Styrene (ABS) using a uPrint SE Plus 3D printer. A DC magnetic field was provided by six NdFeB disk magnets (N42, 6 cm diameter, 5 mm height; Supermagnete) on each side of the animal chamber. As safety precautions, the permanent magnets were covered with a protective lid, and the animal holder base was 3D-printed using the solid option for higher durability. The AC magnetic field was provided by two coils on either side of the animal chamber. A 1 mm thick copper wire was wound around a 3D-printed plastic coil frame with 360 turns each. Corresponding coil-pair resistance was 4.94 Ohm, and coil-pair inductance was 24.5 mH. A Voltcraft 8210 signal generator was used to provide a 140 Hz sine wave, which was amplified using a QSC-GX7 power amplifier. These were then connected to the AC coils. AC and DC magnetic field magnitudes were verified with a magnetometer. For all AC and DC stimulation experiments, mice were stimulated with the coil turned on for 120 s. For DC only stimulation experiments, mice were placed into the animal chamber for 120 s with the coil remaining off.

### Description and timelines of animal experimental procedures

#### Phase I: Toxicity assessment

We first adjusted optimal concentration of MENPs. Three doses were tested, including; 25, 50 and 100 mg/ml. Mice were randomly assigned to either: 25, 50 or 100 mg/ml test groups (n = 8) and received stereotactic injection of MENPs (**Fig. S4A**). Animals were monitored for signs of sub- or epidural hemorrhage, neurological symptoms of the injection, welfare (weight, responsiveness, water intake), discomfort/pain. No animals were eliminated from the experiments due to failing these criteria. Fourteen days after the surgery, mice were sacrificed for immunohistochemical (IHC) analysis of the brain as described below. Brain sections were processed using antibodies raised against astrocytes and microglia (**Fig. S4B**,**C**). Another series of brain sections were stained using standard hematoxylin and eosin (H&E) to evaluate tissue damage at the site of injection (**Fig. S4D**).

#### Phase II: Persistence of nanoparticles at injection site and *c-fos* protein expression

Mice were randomly assigned to three test groups (n = 8) and received stereotactic injection of MENPs (100 mg/ml). We tested the washout of MENPs at different time-points including 48 hours, 2 and 4 weeks (**Fig. S4E**,**F**). At the end of each time-pint, mice underwent transcardial perfusion and brains were removed and used for IHC and H&E analysis. In order to evaluate *c-fos* protein expression, two hours prior to perfusion, half of the mice in each group underwent magnetic stimulation for 120 s. As a control group, the other half of the mice were placed in the coil with no current running through the coil, exposing them only to the DC magnetic field of the permanent magnets.

#### Phase III: Behavioral testing

In order to evaluate the effect of MENP-induced neuronal modulation on brain tissue, two groups of animals were tested and behavioral responses were evaluated. Mice were randomly assigned into two groups (n = 10) and received stereotactic injection of either MENPs or MSNPs (100 mg/ml). Following the recovery period of 1 week post-surgery, animals were stimulated in the magnetic field and behavioral testing was conducted. Specifically, animals were stimulated with either an AC and DC magnetic field (in the *in vivo* coil system with the coil on), or with only a DC magnetic field (in the *in vivo* coil system with the coil off). Measured behavioral parameters were compared between the same mice following stimulation with an AC and DC magnetic field versus stimulation with only a DC magnetic field. At the end of the behavioral testing phase (6 weeks post-surgery), mice underwent transcardial perfusion as described below, and brains were removed and used for IHC analysis (**Fig. S4G**).

### Behavioral testing

#### CatWalk video recording

An automated gait analysis system CatWalk XT (Noldus 7.1, Wageningen, the Netherlands) was to evaluate motor behaviour. The CatWalk consists of an enclosed walkway with a glass plate and a speed video recording camera (**Fig. 3H**). Gait performance was assessed and recorded using the CatWalk analysis software. The glass plate was cleaned and dried before testing each subject to minimize the transmission of olfactory clues and prevent animals from stopping to smell or explore something during a run. In general, one successful test recording consisted of an average of five uninterrupted runs having a comparable running speed with a maximum variation of 30%. The following 20 static and dynamic parameters assessing individual paw functioning and gait patterns were analyzed: stance, mean intensity, print area, print length, print width, swing mean, swing speed, stride length, maximum intensity at maximum contact, maximum intensity, minimum intensity, step cycle, duty cycle, regularity index, base of support of the forelimbs, base of support of the hindlimbs, three limb support, speed, and cadence.

#### Rotarod test

An accelerating rotarod with a grooved rotating beam (3 cm) raised 16 cm above a platform (model 47650, Ugo Basile Biological Research Apparatus, Italy) was used to measure coordination. The latency to fall off the rotating rod was recorded. Data were expressed as the mean value from three trials. Mice were subjected to four 300 s trials per day for three consecutive days (days 1–3) with an inter-trial interval of ∼ 15 min. Mice were forced to run on a rotating drum with speeds starting at 4 rpm and accelerating to 40 rpm within 300 s. Mice remaining on the beam during the full 300 s of the task were taken from the rotarod and given the maximum score.

### Animal sacrifice protocol for immunohistochemical analysis of brain tissue

Mice were deeply anaesthetized with pentobarbital and transcardially perfused with tyrode buffer, followed by ice-cold 4% paraformaldehyde fixative in 0.1 M phosphate buffer. The brains were extracted from the crania and post-fixed in 4% paraformaldehyde overnight, then submerged in sucrose for cryoprotection (24 hours in 20% sucrose at 5°C). Coronal brain sections (20 μm) were cut on a cryostat and stored at −80°C.

### Immunohistochemistry

Tissue sections were incubated overnight with polyclonal rabbit antibodies raised against *c-fos* protein (1:1000; Santa Cruz Biotechnology Inc.; sc-253), GFAP (1:1000; Dako; Z-033429), or Iba-1(1:1000; Wako; 016-26461). *c-fos* IHC used biotinylated donkey anti-rabbit secondary antibody (1:400; Jackson Immunoresearch Laboratories Inc.; 711065152) and avidin–biotin peroxidase complex (1:800, Elite ABC-kit, Vectorlabs; PK-6100). The staining was visualized by 3,3′-Diaminobenzidine (DAB) combined with NiCl2 intensification. GFAP and Iba-1 were visualized using immunofluorescence with donkey anti-rabbit Alexa 488 (1:100; Invitrogen; A-21206). Due to suboptimal perfusion fixation in some animals, brains were not processed for IHC.

### Quantification of *c-fos* immunohistochemically labeled cells

Photographs of the stained motor cortex and thalamus sections from 3 rostrocaudal anatomical levels from bregma (AP −0.58, −0.94 and −1.22) were taken at 10X magnification. We used Cell P software (Olympus Soft Imaging Solutions, Münster, Germany) from an Olympus U-CMAD-2 digital camera connected to an Olympus AX 70 microscope (Olympus, Zoeterwoude, The Netherlands). In the images of the area of interest, the number of c-Fos-positive cells were counted using ImageJ software (version 1.52; NIH, Bethesda, USA). Cells immunopositive for *c-fos* were counted manually, and the mean number of cells was corrected for surface area and expressed as cells/mm^2^. A cell was regarded positive when the intensity of the cell staining was significantly higher than the surrounding background. The average value of three sections was used for statistical analysis in each subject. For the subthalamic nucleus, a digital photograph was taken at one anatomical bregma (−2.06) and all *c-fos* positive cells within 1 mm^2^ of the injection site were counted.

### Quantification of GFAP and Iba-1 immunohistochemically labeled cells

Photographs of the stained motor cortex and thalamus sections from 3 rostrocaudal anatomical levels from bregma (AP −1.70, −2.06 and −2.30) were taken at 10x magnification. We used Cell P software (Olympus Soft Imaging Solutions, Münster, Germany) from an Olympus U-CMAD-2 digital camera connected to an Olympus AX 70 microscope (Olympus, Zoeterwoude, The Netherlands). In the images of the area of interest, fluorescent density was measured using ImageJ software (version 1.52; NIH, Bethesda, USA). The average value of three sections was used for statistical analysis in each subject.

### Statistical analysis

Unless otherwise indicated, data are presented as individual values with bars showing the mean ± standard deviation. The AC magnetic field magnitude and frequency dependence on MENP voltage output was determined using a linear regression, with coefficient of determination presented as R^2^. *In vitro* Ca^2+^ transient activity and *in vivo* cFos expression were analyzed using a one-way analysis of variance (ANOVA) with Tukey’s post-test to compare all groups (**Fig. S4**). *In vitro* analysis of Ca^2+^ signaling with inhibitors was analyzed using a one-way ANOVA with Dunnett’s post-test, using drug-untreated cells as the controls. *c-fos* protein expression in brain tissue was analyzed using an unpaired t-test. Changes in behavioral parameters in the same mice following stimulation with either a DC magnetic field or an AC and DC magnetic field were analyzed using a paired t-test. p-values < 0.05 were considered statistically significant in all cases.

## Acknowledgments

The authors would like to acknowledge Mr. Kerstin Hahn and Prof. Peter van Aken of the Stuttgart Center for Electron Microscopy (StEM, Max Planck Insitute for Solid State Research) for their work capturing TEM images of nanoparticles. We would also like to thank Dr. Samir Hammoud of the Chemical Synthesis facility (Max Planck Insitute for Intelligent Systems) for completing elemental analysis of our nanoparticles. We would also like to thank Mr. Gerd Maier and Dr. Gunther Richter of the Materials Central Scientific Facility (Max Planck Institute for Intelligent Systems) for carrying out XRD analysis of our nanoparticles.

## Author contributions

KK, AJ, MS, and YT conceived the project. KK and AJ designed the experimental layout, analyzed all data, and supervised experiments. KK wrote the manuscript, with the assistance of AJ. YY designed the *in vitro* data acquisition method. KK, YY, and DF carried out *in vitro* modulation experiments. HG conceived and carried out magnetoelectricity measurements. OE and HG designed and fabricated the *in vitro* coil system. OE designed and fabricated the *in vivo* coil system. AJ completed all *in vivo* data acquisition, with the assistance of FA. All authors reviewed and edited the manuscript

## Data and materials availability

All data is available in the main text or the supplementary materials.

## Supplementary Materials

Figures S1-S5

Table S1

Movies S1-S2

