## Supplemental Information for "Injectable Nanoelectrodes Enable Wireless Deep Brain Stimulation of Native Tissue in Freely Moving Mice"

**This PDF file includes:**

Figs. S1 to S5
Table S1
Captions for Movies S1 to S2

**Other Supplementary Materials for this manuscript include the following:**

Movies S1 to S2

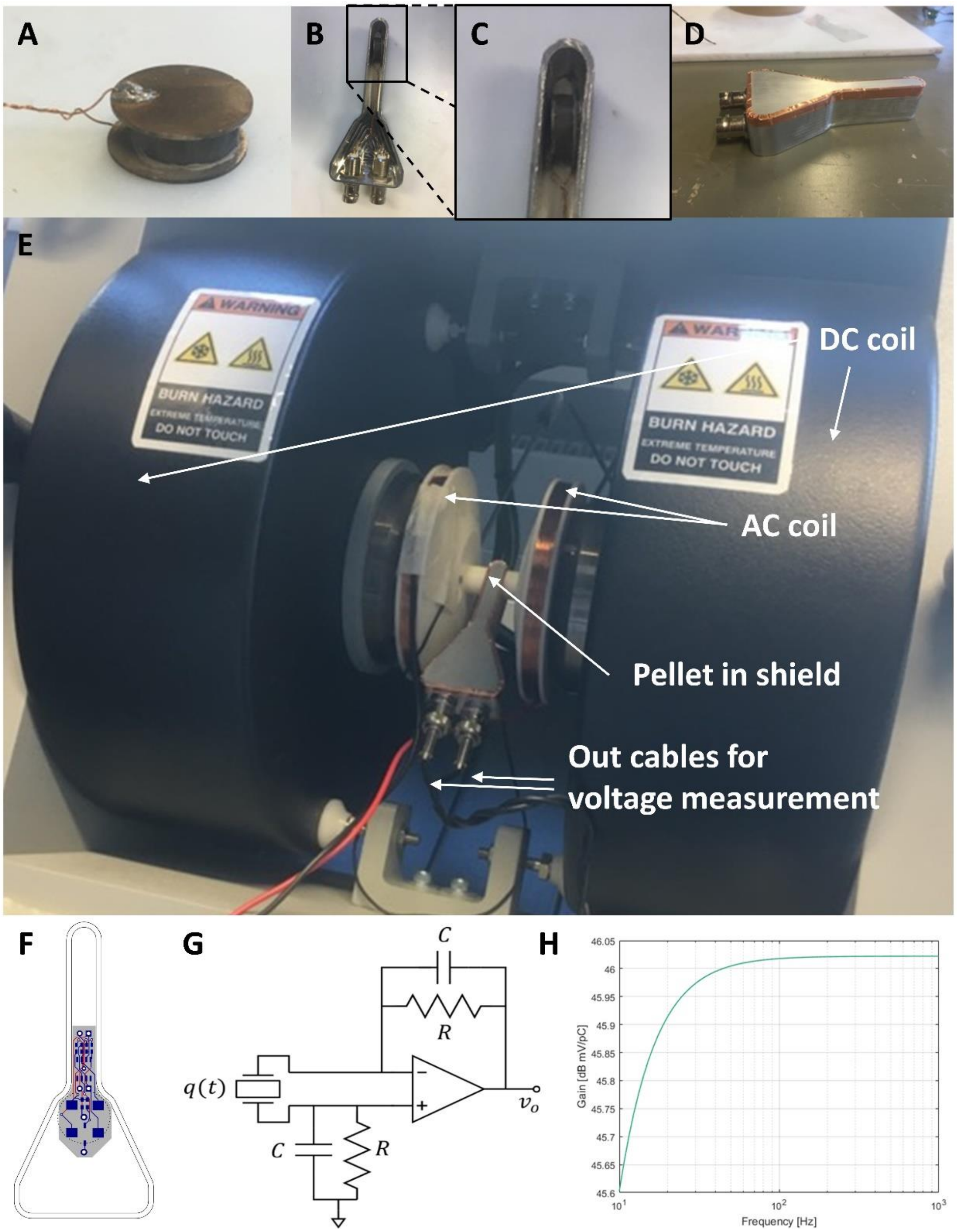


**Figure S1. Magnetoelectricity measurement method.** MENPs (or MSNPs as a control) are pressed into a pellet, and wired to copper plates via conductive paste (A). Pellets are placed into a Faraday shield and wired externally to enable voltage measurement (B). Zoomed image of (B) shows pellet location in Faraday shield (C). The Faraday shield is closed with a lid and conductive copper tape (D). The shield with pellet is placed into a coaxial DC and AC magnetic field and output voltage is measured simultaneously (E). Charge amplifier diagram (F) and circuit schematic (G). Simulated magnitude response function of circuit with $\boldsymbol{R} = \mathbf{5}$ GΩ and $\boldsymbol{C} = \mathbf{10}$ pF (H).

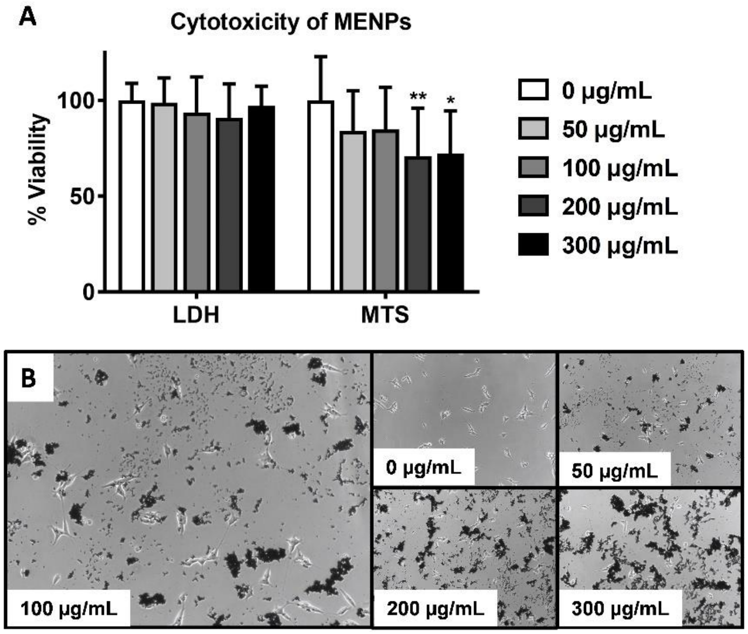


**Figure S2.** ***In vitro*** **toxicity analysis of MENPs.** Quantitative toxicity analysis of MENPs via LDH plasma membrane damage assay and MTS metabolic activity assay (A). Images of cells following 24 h exposure to MENPs at various concentrations. The concentration used in all further *in vitro* experiments was 100 µg/mL (B). Plot shows mean ± SD (n = 3); ANOVA with Dunnett's post-test versus 0 µg/mL MENPs; $**p < 0.01$. $*p<0.05$. unlabeled = ns.

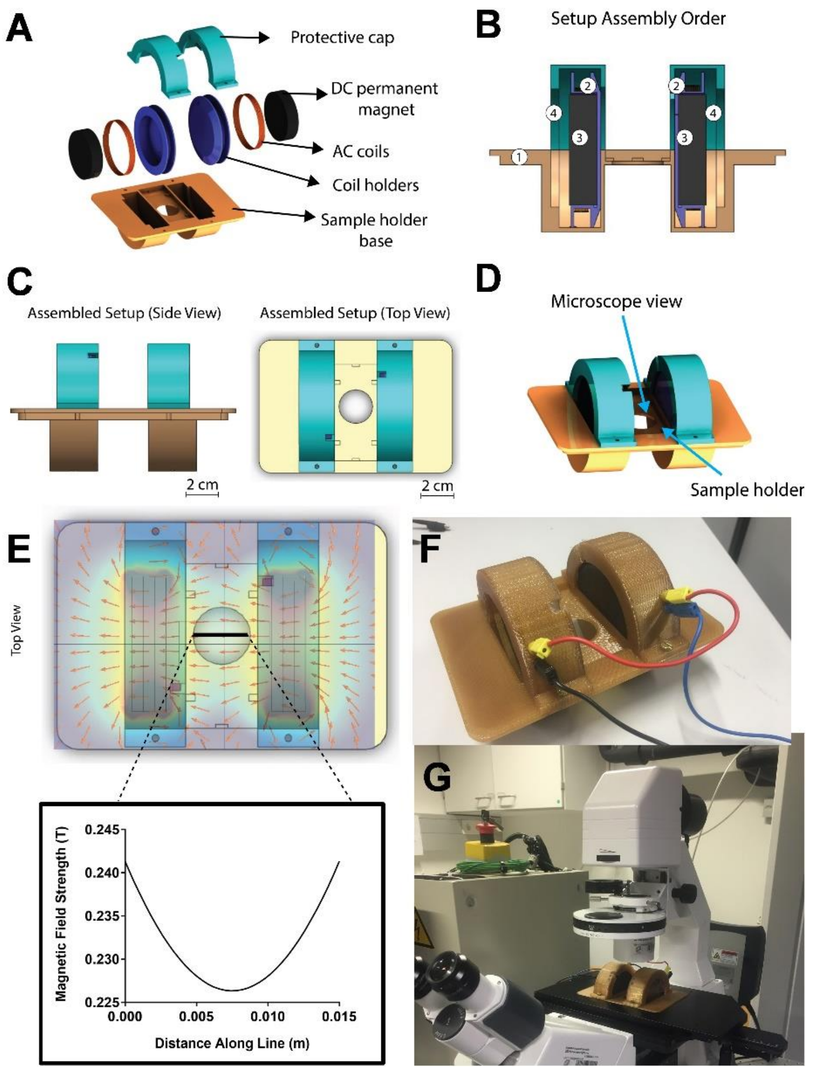


**Figure S3. *In vitro* magnetic stimulation device.** Disassembled schematic shows individual parts of the coil system (A). Internal view of coil system shows layout of components as assembled, with numbers describing the assembly order (B). Schematic of the assembled coil system from top and side views (C) and angled view (D). Plot showing magnitude of the magnetic flux density along the labeled line in the top view in Tesla (E). Picture of the assembled coil (F) and the coil mounted onto the microscope (G).

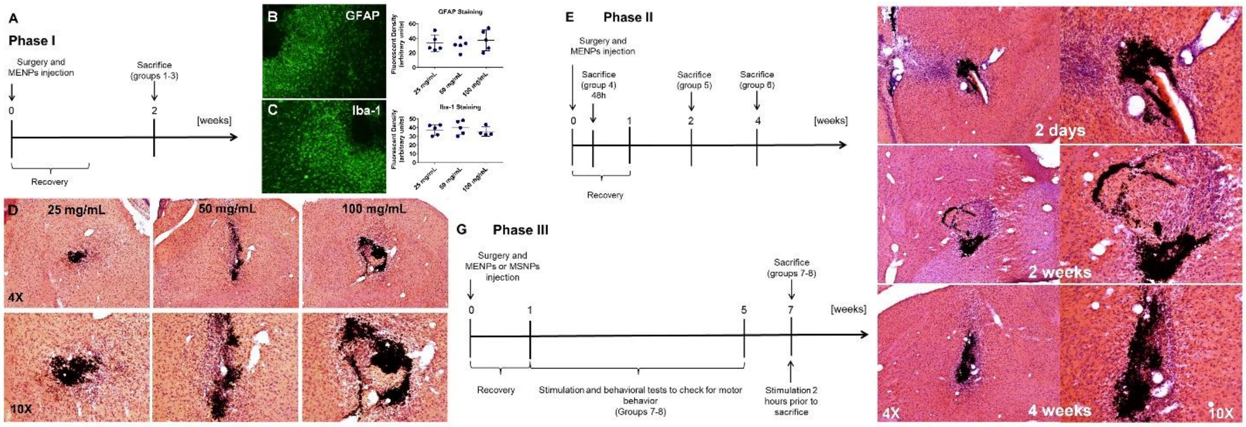


**Figure S4.** ***In vivo*** **experiment layout and analysis of tissue response to MENPs.** Experimental layout of *in vivo* toxicity assessment, Phase 1 (A). GFAP and Iba-1 immunohistochemical staining of tissue surrounding the injection site following injection of 25, 50, or 100 mg/mL MENPs at 2 weeks post-injection (B,C). H&E staining of tissue surrounding injection site following injection of 25, 50, or 100 mg/mL MENPs at 2 weeks post-injection (D). Timeline layout of experimental Phase II (E). H&E staining of tissue surrounding injection site following injection of 100 mg/mL MENPs at 2 days, 2 weeks, and 4 weeks post-injection (F). Timeline layout of experimental Phase III (G).

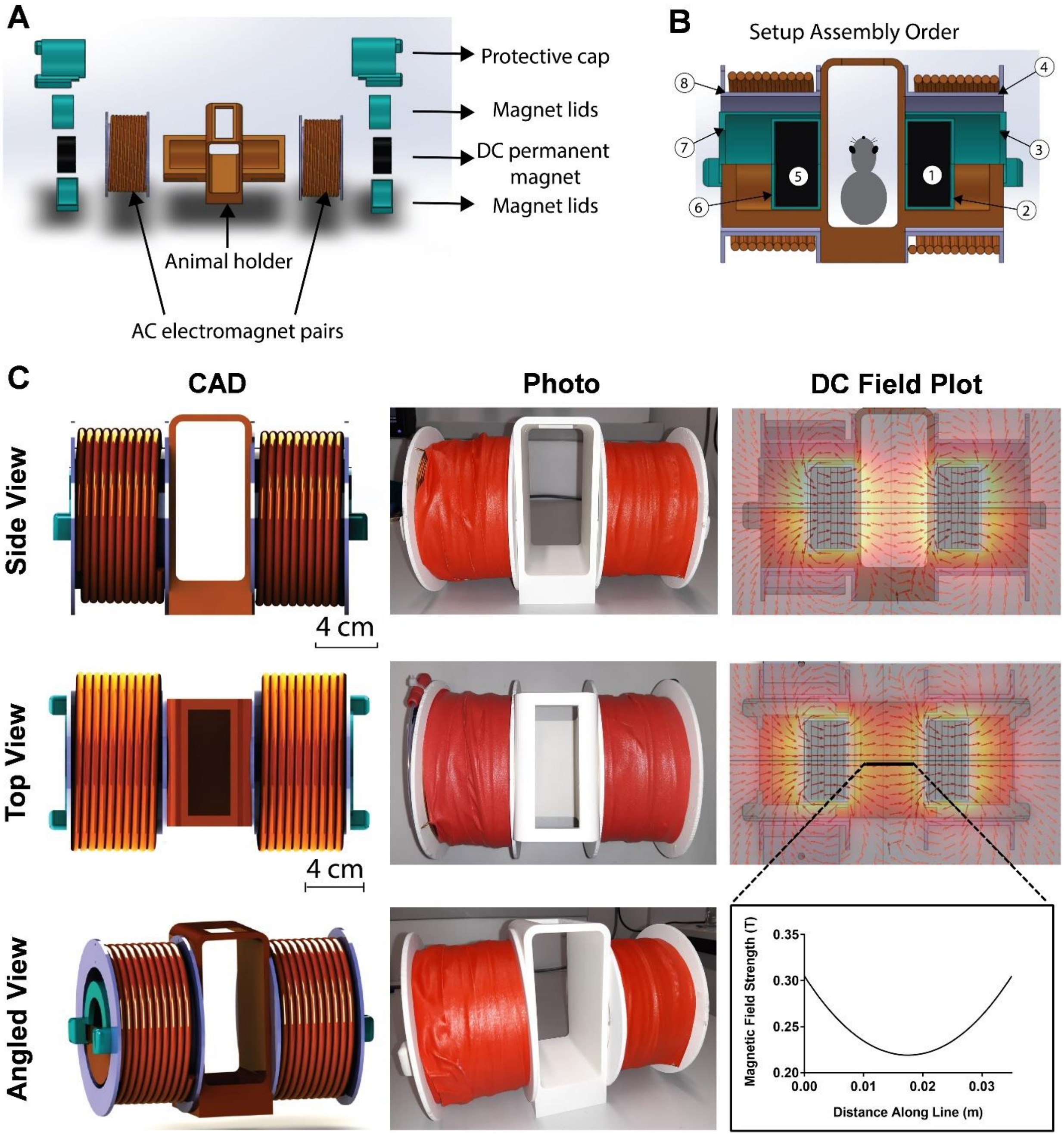


**Figure S5. Schematic and images of *in vivo* magnetic stimulation setup.** Disassembled schematic shows individual parts of the coil system (A). Internal view of coil system shows layout of components as assembled, with numbers describing the assembly order (B). Schematic, picture, and schematic overlaid with DC magnetic field simulation of coil system from the side view and top view (C). Schematic and picture of coil from an angled view. Plot showing magnitude of the magnetic flux density along the labeled line in Tesla (C).

**Table S1.**

p-values following one-way ANOVA with Tukey's post-test for all group comparisons found in **Fig. 2B** are listed, with significant (p < 0.05) values highlighted in yellow (B).

| | No NPs, DC Field | No NPs, AC Field | No NPs, AC+DC Field | MSNPs, No Field | MSNPs, DC Field | MSNPs, AC Field | MSNPs, AC+DC Field | PENPs, No Field | PENPs, DC Field | PENPs, AC Field | PENPs, AC+DC Field | MENPs, No Field | MENPs, DC Field | MENPs, AC Field | MENPs, AC+DC Field |
|---|---|---|---|---|---|---|---|---|---|---|---|---|---|---|---|
| MENPs, AC Field | | | | | | | | | | | | | | | 0.0002 |
| MENPs, DC Field | | | | | | | | | | | | | | >0.9999 | <0.0001 |
| MENPs, No Field | | | | | | | | | | | | | 0.974 | >0.9999 | 0.0013 |
| PENPs, AC+DC Field | | | | | | | | | | | | 0.6824 | >0.9999 | 0.9887 | <0.0001 |
| PENPs, AC Field | | | | | | | | | | | >0.9999 | 0.3883 | 0.9918 | 0.9007 | <0.0001 |
| PENPs, DC Field | | | | | | | | | | 0.9998 | >0.9999 | 0.9608 | >0.9999 | >0.9999 | <0.0001 |
| PENPs, No Field | | | | | | | | | >0.9999 | >0.9999 | >0.9999 | 0.4087 | 0.9975 | 0.9366 | <0.0001 |
| MSNPs, AC+DC Field | | | | | | | | >0.9999 | >0.9999 | >0.9999 | >0.9999 | 0.6869 | >0.9999 | 0.9892 | <0.0001 |
| MSNPs, AC Field | | | | | | | >0.9999 | >0.9999 | >0.9999 | >0.9999 | >0.9999 | 0.5108 | 0.9983 | 0.9543 | <0.0001 |
| MSNPs, DC Field | | | | | | >0.9999 | >0.9999 | >0.9999 | >0.9999 | >0.9999 | >0.9999 | 0.4614 | 0.9967 | 0.9365 | <0.0001 |
| MSNPs, No Field | | | | | >0.9999 | >0.9999 | >0.9999 | >0.9999 | >0.9999 | >0.9999 | >0.9999 | 0.7077 | >0.9999 | 0.9913 | <0.0001 |
| No NPs, AC+DC Field | | | | >0.9999 | >0.9999 | >0.9999 | >0.9999 | >0.9999 | >0.9999 | >0.9999 | >0.9999 | 0.279 | 0.9954 | 0.9007 | <0.0001 |
| No NPs, AC Field | | | >0.9999 | >0.9999 | >0.9999 | >0.9999 | >0.9999 | >0.9999 | >0.9999 | >0.9999 | >0.9999 | 0.844 | >0.9999 | 0.9994 | <0.0001 |
| No NPs, DC Field | | >0.9999 | 0.9417 | 0.9972 | 0.9657 | 0.9773 | 0.9963 | 0.9665 | >0.9999 | 0.9401 | 0.9961 | 0.9987 | >0.9999 | >0.9999 | <0.0001 |
| No NPs, No Field | >0.9999 | >0.9999 | >0.9999 | >0.9999 | >0.9999 | >0.9999 | >0.9999 | >0.9999 | >0.9999 | >0.9999 | >0.9999 | 0.8867 | >0.9999 | 0.9998 | <0.0001 |

**Movie S1.**

This video shows fluctuations in intracellular $Ca^{2+}$ signaling in untreated cells, cells treated with either no NPs, MSNPs, or PENPs and an AC and DC magnetic field, or cells treated with MENPs, and either no field, a DC magnetic field, an AC magnetic field, or an AC and DC magnetic field. (**Fig. 2A-C**). Calibration bars represent $\Delta F/F_o$. Videos are 20X real-time.

**Movie S2.**

This video shows catwalk video recording of mice treated with MSNPs and an AC magnetic field, MSNPs and an AC and DC magnetic field, MENPs and an AC field, or MENPs and an AC and DC magnetic field (**Fig. 3H**).
